# Cytosolic MagLOV Magnetofluorescence in Mammalian Cell Lines and Primary Neurons

**DOI:** 10.64898/2026.09.08.750273

**Authors:** Hengyu Li, Francesco Plastina, Haein An, Ernesto Criado-Hidalgo, Alen Pavlič, Brian L. Zhong, Di Wu, Mikhail G. Shapiro

**Author notes:** Correspondence (D.W.), (M.G.S.).

## Abstract

Magnetic fields can influence the outcome of photochemical reactions through the radical pair mechanism, but whether this sensitivity extends to standard experimental conditions in mammalian cells has remained unclear. Here, we quantify the cytosolic magnetofluorescence of MagLOV – a genetically encoded, flavin-binding fluorescent protein – in mammalian cells, validated against extensive artifact controls. Excitation intensity and magnetic field strength differentially tune response kinetics and amplitude, respectively, with amplitude saturating above approximately 8 mT. Magnetic field-effect amplitude is further modulated by cellular culture state. This response generalizes across HEK293T, HeLa, U2OS, and A549 cells and primary mouse cortical neurons, with plateau amplitudes ranging from 1.4% to 2.7%. Together, these results establish that genetically encoded spin-dependent photochemistry can be quantitatively interrogated under standard mammalian live-cell imaging conditions, and generalizes across mammalian cell types.

---

Photochemical radical pairs provide one of the few established molecular mechanisms by which magnetic fields can alter chemical reaction outcomes. In these systems, the spin states of transiently formed radical pairs govern their probability of recombination versus separation^1–3^. This mechanism has been most extensively studied in the context of animal magnetoreception, where flavin-based photoreceptors of the cryptochrome family have been proposed to sense the geomagnetic field through spin-correlated radical pairs formed between a flavin cofactor and a nearby tryptophan residue^4–8^. More broadly, magnetic field effects (MFEs) on flavin photochemistry raise the possibility that spin-dependent chemistry is not restricted to specialized magnetoreceptor proteins, but may be a more general feature of flavin-binding photoreceptor domains^9–19^.

Consistent with this possibility, the light-oxygen-voltage (LOV) domain family, structurally and photochemically distinct from cryptochromes but similarly reliant on flavin photochemistry, has recently been found to exhibit robust magnetic field-dependent fluorescence^20–22^. Directed evolution of the AsLOV2 domain^23^ yielded MagLOV, a genetically encoded, cofactor-independent fluorescent protein whose fluorescence is modulated by external magnetic fields with substantially larger amplitude than its parental domain^21^. Because MagLOV requires no exogenous cofactor and remains magnetosensitive in bacterial cells and in purified form, it offers a tractable, genetically encoded system for studying spin-dependent photochemistry directly in living cells.

Beyond its use as a fluorescent reporter, MagLOV has recently been engineered through directed evolution to enable optically detected magnetic resonance (ODMR) in living bacterial cells, extending genetically encoded magnetofluorescence toward quantum-sensing applications such as radiofrequency detection and fluorescence-based magnetic resonance imaging^24^. This advance is part of a broader, largely concurrent body of work establishing optically addressable spin states in flavin-associated fluorescent proteins, including radiofrequency-controlled spin-correlated radical pair dynamics in a red-fluorescent-protein-flavin system in a living transgenic animal^25,26^, optically detected magnetic resonance in purified cryptochrome and the LOV-family fluorescent reporter iLOV^27,28^, and coherent optical spin control of a mechanistically distinct fluorescent-protein qubit EYFP^29^. Collectively, these advances establish flavin- and fluorescent-protein-based systems as a versatile platform for engineering spin-dependent readouts with tunable magnetic field sensitivity and resonance properties.

Native cytosolic MagLOV, expressed at room temperature in living mammalian cells, has appeared in the literature only anecdotally or as a proof-of-imaging feasibility. The original report of MagLOV noted magnetoresponsiveness in mammalian cells without accompanying quantitative data^21^, and a subsequent live-cell quantum-imaging platform confirmed that MagLOV can be expressed and imaged in HeLa cells but did not characterize its magnetic field response^30^. Quantitative MFE data for MagLOV-family proteins have instead been obtained in settings distinct from this one. In Escherichia coli at room temperature, MFEs have been characterized for engineered MagLOV2^31^ and, at high magnetic field strengths, for the wild-type, unevolved AsLOV2 domain in vitro^32^. Directed evolution of MagLOV2 has further enabled ODMR and quantum-sensing applications, again in bacterial cells^24^, and a mitochondria-targeted MagLOV2 variant has been characterized in mammalian cells^33^. Comparable magnetic field sensitivity has also been documented in mechanistically distinct fluorescent-protein-flavin systems, including a red fluorescent protein studied in vitro^34^ and in a transgenic nematode^25^, and in a fluorescent-protein spin qubit, EYFP, in mammalian cells cooled to cryogenic temperature^29^. Whether the native, cytosolic MagLOV response itself is robust, quantitatively tunable by physical inputs, and generalizable across mammalian cell types under standard live-cell imaging conditions has not been established.

Here, we address these questions directly. We establish a quantitative assay for cytosolic MagLOV magnetofluorescence in mammalian cells, validated against extensive artifact controls. Using this assay, we show that excitation intensity and magnetic field strength differentially tune response kinetics and amplitude, respectively. We further find that MFE amplitude is modulated by cellular culture state. Finally, we demonstrate that cytosolic MagLOV magnetofluorescence generalizes across HEK293T, HeLa, U2OS, and A549 cells and primary mouse cortical neurons. Together, these results establish that genetically encoded spin-dependent photochemistry can be quantitatively interrogated in mammalian cells, where its behavior is jointly shaped by physical inputs and cellular state.

## RESULTS

### Robust and reversible MagLOV magnetofluorescence in the mammalian cytosol

To establish a quantitative assay for MagLOV magnetofluorescence in the mammalian cytosol, we expressed MagLOV in HEK293T cells and monitored its fluorescence while periodically switching an external magnetic field on and off (**Fig. 1a**). Cells were subjected to a 60 mT magnetic field in alternating 10-s OFF and ON intervals during optical excitation. Magnetic field application produced a readily detectable decrease in MagLOV fluorescence that reversed upon field removal and was reproducible over successive switching cycles (**Fig. 1b–d**). In contrast, identical servo movement in the absence of the magnet produced minimal phase-locked fluorescence changes, excluding mechanical perturbation of the imaging setup as the dominant source of the response (**Fig. 1d**). Single-cell quantification of the raw fluorescence decrease between individual field-off and field-on images likewise revealed a readily detectable response (**Fig. 1c**), with the amplitude of this single-timepoint comparison independently quantified as a drift-corrected plateau value across replicates below (**Fig. 1e**).

**Figure 1.**
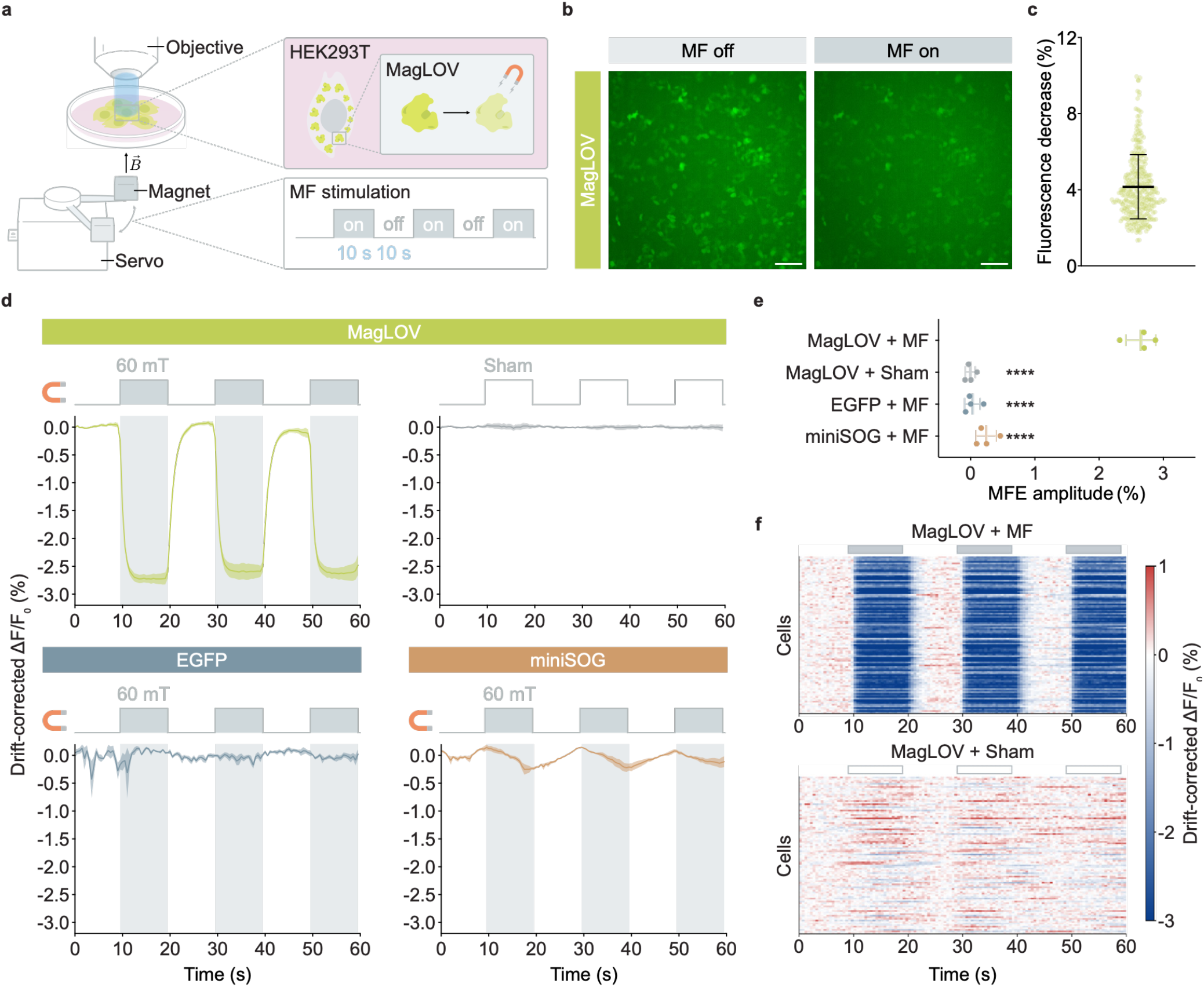
Robust and reversible MagLOV magnetofluorescence in the mammalian cytosol. **a**, Schematic of fluorescence imaging of HEK293T cells expressing cytosolic MagLOV during periodic magnetic-field stimulation. A servo-driven permanent magnet alternated the field between OFF and ON states at 10-s intervals during optical excitation. **b**, Representative fluorescence images of MagLOV-expressing HEK293T cells acquired during magnetic-field-off and magnetic-field-on periods. Images are displayed using identical intensity scaling. Scale bar, 100 µm. **c**, Single-cell fluorescence decrease between the field-OFF and field-ON images shown in **b.** Each point represents one cell (n = 332 cells); line and error bars indicate the mean ± s.d. **d**, Drift-corrected fluorescence responses of cells expressing MagLOV during 60 mT magnetic-field stimulation or sham servo movement, and of cells expressing EGFP or miniSOG during 60 mT stimulation. Filled and outlined regions denote field-ON and sham-servo phases, respectively. Lines and envelopes indicate mean ± s.e.m. across four independent biological replicates. **e**, Plateau MFE amplitudes calculated from drift-corrected single-cell fluorescence traces (see Methods). Each point represents one independent biological replicate; lines and error bars indicate mean ± s.d. (n = 4). Statistical significance was determined by one-way ANOVA followed by Dunnett’s multiple-comparisons test using MagLOV + MF as the reference (****P < 0.0001 for all comparisons). **f**, Single-cell heat maps of drift-corrected MagLOV responses during magnetic-field stimulation or sham servo movement. One hundred cells are shown per condition (25 randomly sampled from each of four biological replicates), intermixed without sorting by response magnitude. Identical linear color scales were used for both conditions.

To distinguish the MagLOV response from nonspecific fluorescence changes during illumination, we performed the same measurements in cells expressing EGFP or another flavin-binding protein, miniSOG^35^. EGFP showed no appreciable field-locked fluorescence response, while miniSOG exhibited a smaller but detectable response (**Fig. 1d**). Quantification across four independent biological replicates yielded a plateau MFE amplitude of approximately 2.6% for MagLOV, substantially exceeding the responses observed with sham stimulation, EGFP, and miniSOG (**Fig. 1e**). The small miniSOG response suggests that magnetic sensitivity is not unique to MagLOV, although its magnitude appears strongly dependent on protein context. At the single-cell level, repeated field-locked fluorescence decreases were observed across the MagLOV-expressing population, whereas cells subjected to sham stimulation lacked this response (**Fig. 1f**).

Because prolonged optical excitation introduced slow baseline drift, we corrected fluorescence traces using a log-space regression model that isolated the phase-dependent signal from nuisance drift and startup transients (**Extended Data Fig. 1b**), and next tested whether the observed MFE was robust to this correction and analysis procedure. Repeated MagLOV responses remained evident in baseline-normalized traces before correction, while direction-specific analysis of the raw fluorescence preserved the field-associated response and largely cancelled drift-associated contributions in the control conditions (**Extended Data Fig. 1a,c**). As an independent, correction-free robustness analysis, a direction-balanced estimator calculated directly from raw fluorescence remained positive for MagLOV across a range of post-switch analysis windows, whereas the control conditions remained near zero (**Extended Data Fig. 1d**). Together, these analyses show that the field-locked MagLOV response is not dependent on a particular drift-correction or quantification procedure and establish a robust and reversible cytosolic MFE suitable for further quantitative characterization.

### Excitation intensity and magnetic-field strength differentially tune MagLOV amplitude and kinetics

With a quantitative cytosolic MagLOV assay established, we next examined how excitation intensity and magnetic field strength shape response amplitude and dynamics. We first varied excitation intensity from 1.65 to 82.5 mW cm^-2^ while maintaining the magnetic field at 60 mT (**Fig. 2a**). Reversible fluorescence responses were observed across the full range of excitation intensities, but their amplitude and kinetics showed distinct dependencies on illumination. The plateau MFE amplitude was already approximately 3.1% at 1.65 mW cm^-2^, remained near 3.3–3.6% over a broad intermediate range of excitation intensities, and decreased to approximately 2.4% near 82.5 mW cm^-2^ (**Fig. 2c**). In contrast, the normalized ON-response half-time (*t*_1/2_) decreased progressively from approximately 1.7 s at 1.65 mW cm^-2^ to approximately 0.7 s at 82.5 mW cm^-2^ (**Fig. 2c**). Thus, increasing excitation intensity progressively accelerated the MagLOV response, whereas response amplitude varied more weakly over the intermediate illumination range and declined at the highest intensity tested.

**Figure 2.**
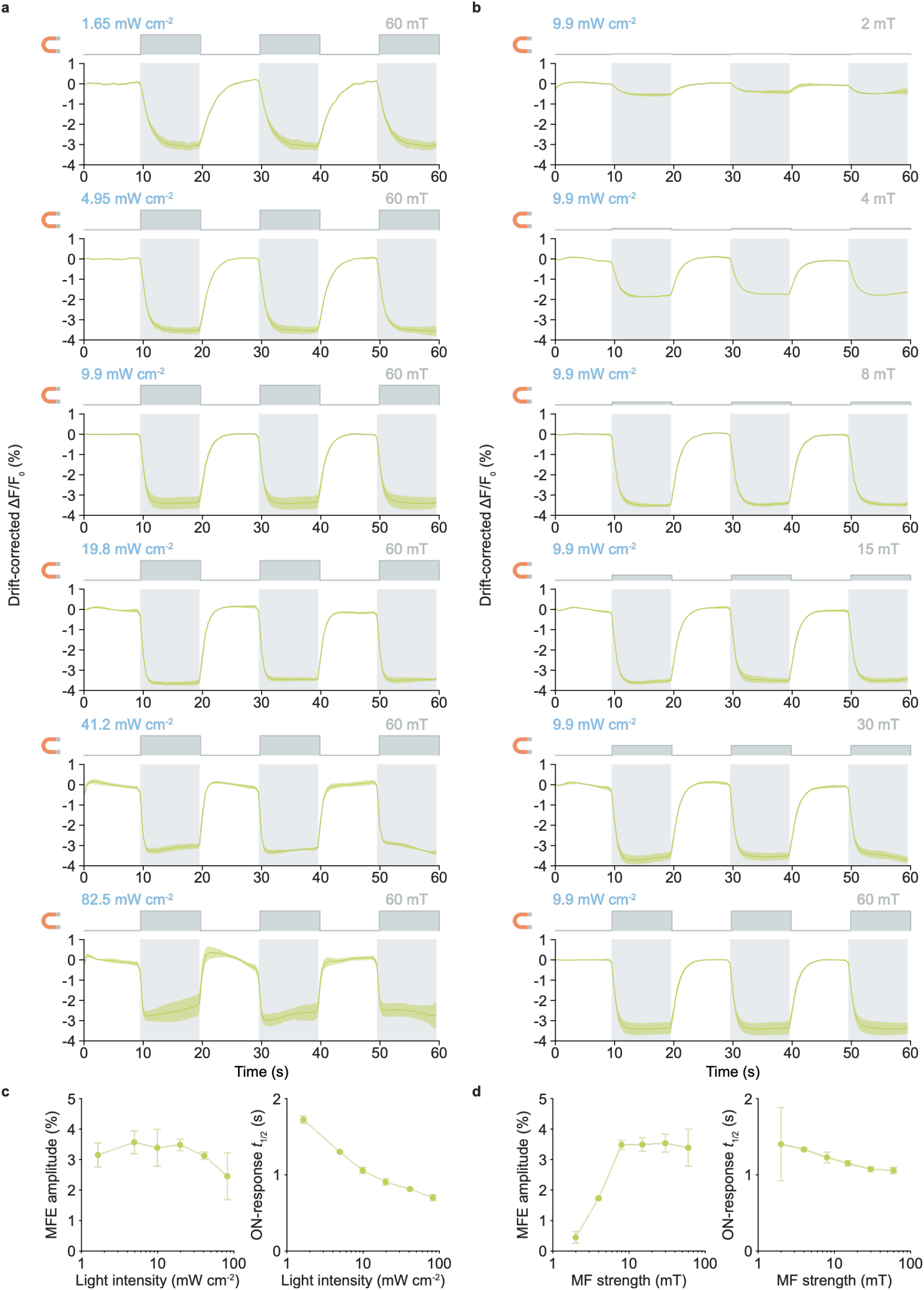
Excitation intensity and magnetic-field strength differentially tune cytosolic MagLOV amplitude and kinetics. **a**, Drift-corrected fluorescence responses of MagLOV-expressing HEK293T cells during repeated 60 mT magnetic-field stimulation at the indicated excitation intensities (1.65–82.5 mW cm^-2^). Shaded regions indicate magnetic-field-on intervals. Lines and envelopes indicate the mean ± s.e.m. across independent biological replicates. **b**, Drift-corrected fluorescence responses during stimulation with the indicated magnetic-field strengths (2–60 mT) at a fixed excitation intensity of 9.9 mW cm^-2^. Shaded regions indicate magnetic-field-on intervals. Lines and envelopes indicate the mean ± s.e.m. across independent biological replicates. **c**, Quantification of excitation-intensity dependence at a fixed magnetic-field strength of 60 mT. Left, plateau MFE amplitude calculated from drift-corrected single-cell traces (see Methods). Right, normalized ON-response half-time (*t*_1/2_) (see Methods). **d**, Quantification of magnetic-field-strength dependence at a fixed excitation intensity of 9.9 mW cm^−2^. Left, plateau MFE amplitude calculated as in **c.** Right, normalized *t*_1/2_ as calculated in **c**. In **c** and **d**, individual points represent independent biological replicates, connected symbols indicate the mean, and error bars indicate the s.d. (n = 3–4 independent biological replicates per condition).

We next varied magnetic field strength from 2 to 60 mT while maintaining the excitation intensity at 9.9 mW cm^-2^ (**Fig. 2b**). The plateau MFE amplitude increased steeply over the lower part of this range, from approximately 0.4% at 2 mT to approximately 1.7% at 4 mT and approximately 3.4% at 8 mT, after which little additional increase was observed up to 60 mT (**Fig. 2d**). Normalized response kinetics also accelerated with increasing field strength, but this dependence was comparably modest, with *t*_1/2_ decreasing from approximately 1.4–1.5 s at 2 mT to approximately 1.1 s at 30–60 mT (**Fig. 2d**). Thus, response amplitude exhibited a pronounced field-dependent increase followed by saturation above approximately 8 mT, whereas response kinetics changed more gradually across the same range. This saturating dose-response profile is characteristic of radical-pair-mediated MFEs^2,3,10^.

Baseline-normalized traces from individual biological replicates reproduced the strong increase in response magnitude between 2 and 8 mT and the limited additional increase at higher field strengths (**Extended Data Fig. 2b**). Increasing excitation intensity was accompanied by greater baseline drift, particularly at the highest intensity tested, although repeated field-locked fluorescence changes remained evident in the uncorrected traces (**Extended Data Fig. 2a**). The reduction in plateau MFE amplitude at 82.5 mW cm^-2^ therefore occurred under the condition with the greatest illumination-associated baseline instability and should be interpreted in this context. Together, these measurements show that excitation intensity and magnetic field strength differentially shape cytosolic MagLOV responses. Field strength primarily controls response magnitude over the lower-field regime before saturation, whereas excitation intensity has a particularly strong influence on response kinetics.

### Cellular culture state tunes MagLOV magnetofluorescence

Having established how physical inputs shape the cytosolic MagLOV response, we next asked whether cellular state also modulates this behavior, given that cellular physiology is known to vary substantially with culture conditions. To vary cellular state independently of excitation intensity and magnetic field strength, we exposed HEK293T cells to fresh culture medium for defined durations (4–20 h) before imaging, while holding excitation intensity (9.9 mW cm^-2^) and magnetic field strength (4 mT) constant (**Fig. 3a**).

**Figure 3.**
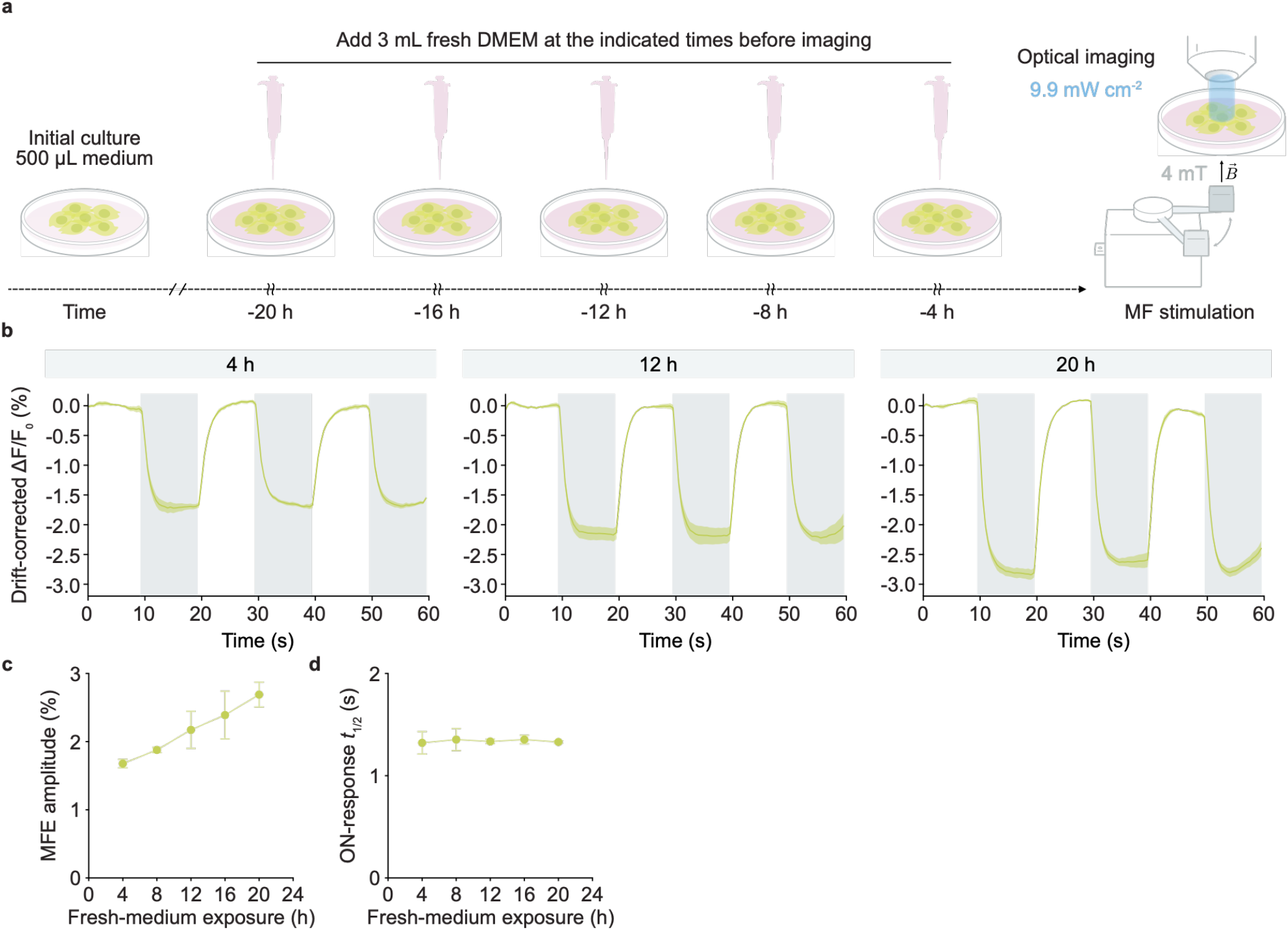
Cellular culture state tunes MagLOV magnetofluorescence. **a**, Schematic of the fresh-medium exposure protocol. HEK293T cells were transfected and maintained in 500 μL medium, with 3 mL fresh DMEM added at the indicated times before a common imaging endpoint (see Methods). Imaging was performed at an excitation intensity of 9.9 mW cm^-^ 2 and a magnetic-field strength of 4 mT. **b**, Drift-corrected fluorescence responses of MagLOV-expressing HEK293T cells during repeated 4-mT magnetic-field stimulation following the indicated durations of fresh-medium exposure (4, 12, or 20 h; representative of the full range shown in **c** and **d**). Shaded regions indicate magnetic-field-on intervals. Lines and envelopes indicate the mean ± s.e.m. across four independent biological replicates (n = 4). **c**, Quantification of the fresh-medium exposure-time dependence of plateau MFE amplitude, calculated from drift-corrected single-cell traces (see Methods). Points and error bars indicate the mean ± s.d. across four independent biological replicates (n = 4). **d**, Quantification of the fresh-medium exposure-time dependence of normalized ON-response *t*_1/2_, calculated as in **Fig. 2c** (see Methods). Points and error bars indicate the mean ± s.d. across four independent biological replicates (n = 4).

Longer fresh-medium exposure progressively increased the plateau MFE amplitude, from approximately 1.7% at 4 h to approximately 2.7% at 20 h (**Fig. 3b,c**), without a corresponding change in response kinetics. The normalized ON-response *t*_1/2_ remained essentially constant (approximately 1.3–1.4 s) across the full range of fresh-medium exposure times tested (**Fig. 3d**), indicating that cellular state selectively tunes response amplitude while leaving response kinetics largely unaffected. Baseline-normalized traces from individual biological replicates reproduced this trend prior to drift correction (**Extended Data Fig. 3a**).

Because prolonged culture without fresh medium is associated with changes in basal fluorescence, we next examined whether basal MagLOV fluorescence itself varied with fresh-medium exposure. Basal fluorescence increased modestly with longer exposure, rising to approximately 112% of its 4-h value by 20 h (**Extended Data Fig. 3b**), whereas MFE amplitude rose to approximately 160% of its 4-h value over the same range (**Extended Data Fig. 3c**), which is a substantially larger relative change than that observed for basal fluorescence. Across all biological replicates, MFE amplitude was positively correlated with basal fluorescence (Pearson r = 0.64, P = 0.002; **Extended Data Fig. 3d**). However, because both basal fluorescence and MFE amplitude independently increased with fresh-medium exposure time, this correlation could reflect a shared dependence on exposure time rather than a direct relationship between fluorescence level and response amplitude. To distinguish between these possibilities, we centered basal fluorescence and MFE amplitude within each exposure-time group before pooling replicates for correlation analysis. This centered correlation was not significant (Pearson r = -0.16, P = 0.491; **Extended Data Fig. 3e**), indicating that the raw correlation between basal fluorescence and MFE amplitude is driven by their shared dependence on fresh-medium exposure time, rather than reflecting a direct relationship between the two variables. Together, these results show that cytosolic MagLOV magnetofluorescence is tunable by cellular culture state, independently of the physical inputs characterized above.

### MagLOV magnetofluorescence is reproducible across diverse mammalian cellular backgrounds, including primary neurons

To determine whether cytosolic MagLOV magnetofluorescence extends beyond HEK293T cells, we examined the response in three additional mammalian cell lines, HeLa, U2OS, and A549, as well as primary mouse cortical neurons. MagLOV was expressed in the cytosol and imaged during repeated 60 mT magnetic field stimulation at an excitation intensity of 1.65 mW cm^-2^ (**Fig. 4a**). In all four cellular backgrounds, magnetic field application produced a reversible decrease in MagLOV fluorescence that recovered upon field removal and was reproduced over successive stimulation cycles (**Fig. 4b**). Quantification across independent biological replicates yielded plateau MFE amplitudes of approximately 1.9% in HeLa, 2.2% in U2OS, 1.4% in A549, and 1.8% in primary neurons (**Fig. 4c**). Thus, readily detectable cytosolic MagLOV magnetofluorescence is preserved across distinct mammalian cellular backgrounds, including primary neurons (n = 4 independent biological replicates per cellular background).

**Figure 4.**
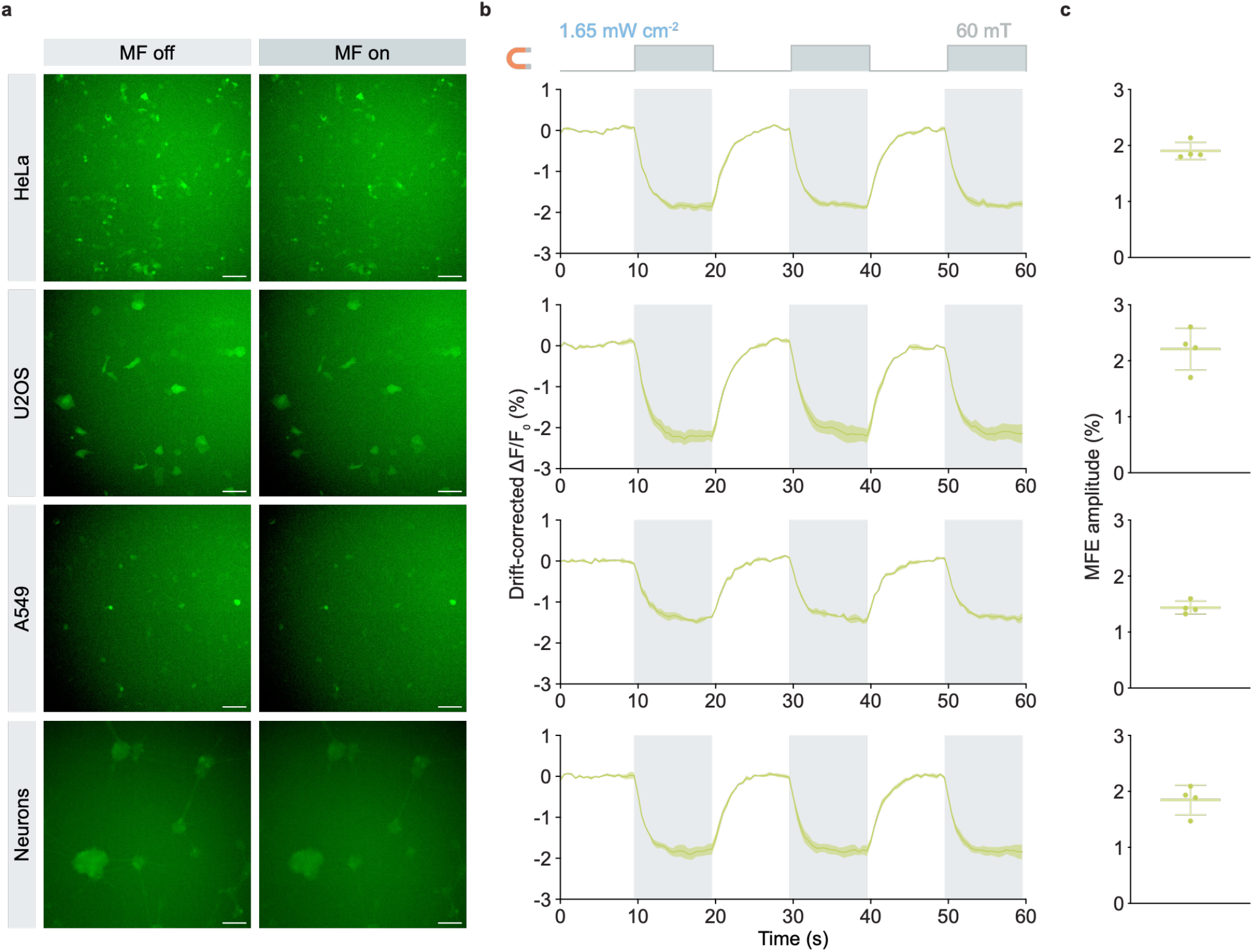
MagLOV magnetofluorescence is reproducible across diverse mammalian cellular backgrounds, including primary neurons. **a**, Representative fluorescence images of MagLOV-expressing HeLa, U2OS, and A549 cells and primary mouse cortical neurons acquired during magnetic-field-off (MF off) and magnetic-field-on (MF on) periods. The same linear brightness and contrast settings were applied to the MF-off and MF-on images within each cellular background. The three tumor cell lines were displayed using a common intensity range, whereas primary-neuron images were displayed using a separate range because they were acquired in an independent experiment. Display settings are not intended for quantitative comparison of absolute fluorescence across cellular backgrounds. Scale bars, 100 µm. **b**, Drift-corrected fluorescence responses of MagLOV-expressing HeLa, U2OS, and A549 cells and primary mouse cortical neurons during repeated 60 mT magnetic-field stimulation at an excitation intensity of 1.65 mW cm^-2^. Shaded regions indicate magnetic-field-on intervals. Lines and shaded envelopes indicate the mean ± s.e.m. across four independent biological replicates for each cellular background. **c**, Plateau MFE amplitudes calculated from drift-corrected single-cell fluorescence traces (see Methods). Each point represents one independent biological replicate. Horizontal lines and error bars indicate the mean ± s.d. (n = 4 independent biological replicates per cellular background).

We next examined the robustness of these responses at the biological-replicate and stimulation-cycle levels. Field-locked fluorescence decreases remained evident in baseline-normalized traces from individual biological replicates before drift correction, despite differences in slow baseline behavior among cellular backgrounds (**Extended Data Fig. 4a**). Moreover, cycle-resolved plateau MFE amplitudes showed no significant change across the three successive magnetic field-on intervals in any of the four cellular backgrounds tested (**Extended Data Fig. 4b**). Together with the HEK293T measurements, these results demonstrate that reversible cytosolic MagLOV magnetofluorescence can be reproducibly detected across diverse mammalian cellular contexts and remains stable over repeated magnetic field stimulation.

## DISCUSSION

This study rigorously demonstrates and quantifies cytosolic MagLOV magnetofluorescence in mammalian cells, defining how excitation intensity, magnetic field strength, and cellular state jointly shape its amplitude and kinetics, and demonstrating that this response generalizes across multiple mammalian cell types, including primary neurons. These results show that genetically encoded spin-dependent photochemistry can be elicited and quantitatively interrogated under standard live-cell imaging conditions, providing an experimental foundation for future mechanistic studies and for applications of genetically encoded fluorescent proteins in quantum-sensing contexts^36^.

The differential tuning of response kinetics and amplitude by excitation intensity and magnetic field strength, respectively, indicates that these two physical inputs act on at least partially separable aspects of the underlying photochemistry^2,3,10,37^. This separation offers a practical guide for using MagLOV as a fluorescent reporter: excitation intensity can be adjusted to optimize acquisition speed without substantially compromising MFE amplitude over an intermediate range, whereas magnetic field strength must be maintained above the saturating regime identified here (approximately 8 mT) to achieve near-maximal signal. More broadly, the differential dependence of response kinetics and amplitude on optical and magnetic inputs may provide orthogonal parameters for future spatial encoding strategies.

The generalization of this response across HEK293T, HeLa, U2OS, and A549 cells and primary cortical neurons indicates that cytosolic MagLOV magnetofluorescence is not restricted to a single cellular context, supporting its broader use as a reporter of intracellular spin-dependent photochemistry. Because expression level was not independently quantified across cellular backgrounds in this study, potential contributions of differences in transfection efficiency or protein expression to the observed amplitudes cannot be excluded.

Beyond this comparison across cellular contexts, these results complement a concurrent and rapidly expanding body of work applying genetically encoded, flavin-associated spin systems toward quantum-sensing applications, including engineered MagLOV2 variants for ODMR in bacterial cells, radiofrequency-controlled radical-pair dynamics in a red-fluorescent-protein-flavin system in a living transgenic animal, ODMR in purified cryptochrome and iLOV, and coherent spin control of a mechanistically distinct fluorescent-protein qubit, EYFP. Whereas these studies establish the capabilities of engineered or purified systems under specialized conditions, the present work defines how the native, unengineered MagLOV response behaves under the physical and physiological conditions of routine live-cell fluorescence imaging. The quantitative operating principles established here may in turn inform the design and interpretation of engineered magnetofluorescent variants intended for use in mammalian cells.

This study has several limitations. The biochemical origin of the cytosolic MagLOV response, including the intracellular factors that shape radical-pair yield, has not been fully established in this or prior work, and will require further biochemical and spectroscopic investigation, analogous to approaches used to dissect electron-transfer distance and kinetics in de novo engineered flavin-tryptophan radical pair systems^38,39^. Notably, EGFP itself possesses a well-characterized long-lived triplet state that underlies its own oxidative photochemistry40, yet showed no detectable field-locked response in this study (**Fig. 1d**), underscoring that not all photoactive dark-state chemistry is magnetically responsive under these conditions. The present characterization is also limited to a small number of cell lines and a single type of primary cell; generalization to additional cellular contexts remains to be tested.

Together, these findings demonstrate a robust mammalian cytosolic MFE for MagLOV and establish a quantitative approach to study MagLOV as a reporter of intracellular spin-dependent photochemistry in mammalian cells. Future efforts can build on this foundation to engineer and apply MagLOV and similar proteins toward basic research and potential translational uses.

## ACKNOWLEDGEMENTS

The authors thank A. E. Cohen and K. M. Xiang for scientific discussion, M. Barbic for important early discussions and guidance on the design of the magnetic stimulation setup, A. G. York for sharing the pRSET-MagLOV construct, and A. Yadav, N. N. Nyström, and D. D. Tang for technical support in primary cell culture.

## AUTHOR CONTRIBUTIONS

H.L., D.W., and M.G.S. conceived and planned the study. H.L. designed and performed the experiments and analyzed the data. F.P. contributed to mammalian cell experiments, data processing, and pilot experiments that informed subsequent measurements. H.A. isolated primary mouse cortical neurons. E.C.-H. provided technical assistance with 3D printing of custom experimental components. A.P. contributed acquisition software and thermal-control hardware that were adapted for this study. B.L.Z. provided technical input on the Arduino- and servo-based magnetic stimulation system. H.L. and M.G.S. wrote the manuscript with input from all other authors. D.W. and M.G.S. supervised the research.

## FUNDING

This research was supported by the National Institute of Health (DP1 EB033154 to M.G.S.). E.C.-H. was supported by the James Boswell Postdoctoral Fellowship program between Caltech and Huntington Medical Research Institutes. A.P. was supported by the Swiss National Science Foundation and the Human Frontier Science Program. F.P. was supported by the Summer Undergraduate Research Fellowship program at Caltech.

M.G.S. is an investigator of the Howard Hughes Medical Institute. We apologize for any unintentional omission of relevant citations.

