## Supplementary Information for "Cytosolic MagLOV Magnetofluorescence in Mammalian Cell Lines and Primary Neurons"

### Methods

#### Plasmid construction

The MagLOV coding sequence was PCR-amplified from plasmid pRSET-MagLOV (Addgene plasmid #219957) using Q5 High-Fidelity DNA Polymerase (New England Biolabs) and cloned into the pCMVSp mammalian expression backbone, available in the laboratory, by Gibson assembly[1]. For lentiviral expression, the MagLOV coding sequence was similarly cloned into the pLenti-EF1s-WPRE backbone, also available in the laboratory, to generate pLenti-EF1s-MagLOV-WPRE. The EGFP coding sequence was PCR-amplified from a plasmid available in the laboratory, and the miniSOG coding sequence, reverse-translated from the amino acid sequence reported in ref. 35, was synthesized as a gBlock (Integrated DNA Technologies); both were cloned into the pCMVSp backbone by Gibson assembly. The resulting plasmids were verified by whole-plasmid sequencing (Quintara Biosciences).

#### Mammalian cell culture and transient transfection

HEK293T, HeLa, U2OS, and A549 (ATCC) cells were maintained in DMEM supplemented with 10% fetal bovine serum and 1× penicillin-streptomycin at 37˚C and 5% $CO_2$. For imaging experiments, 14-mm glass-bottom dishes (Avantor) were precoated with bovine collagen (CELLINK) at 100 µg $mL^{-1}$ in 10 mM HCl for 3 h at 37˚C and rinsed with PBS before cell seeding. Cells were seeded 24 h before transfection at densities of $1.6 \times 10^5$ HEK293T, $1.1 \times 10^5$ HeLa, $0.7 \times 10^5$ U2OS, or $1.1 \times 10^5$ A549 cells per dish in 200 µL culture medium. Immediately before transfection, 300 µL fresh medium was added to each dish. Cells were transfected with 320 ng plasmid DNA using PEI-MAX[2] (Polysciences) at a PEI:DNA mass ratio of 2.58:1. The same transfection procedure was used for MagLOV, EGFP, and miniSOG constructs. For the experiments shown in Figs. 1, 2, and 4, cells were imaged 24 h after transfection, with 3 mL fresh medium added 4 h before imaging. For the fresh-medium exposure experiments in Fig. 3, cells were imaged 48 h after transfection, and 3 mL fresh medium was added at the indicated times before imaging.

#### Primary embryonic mouse neuron isolation and culture

All animal procedures were approved by the Institutional Animal Care and Use Committee of the California Institute of Technology. Primary neurons were isolated from embryonic day 17 (E17) mice (Charles River Laboratories). Culture substrates were coated with poly-D-lysine at 100 µg $mL^{-1}$ at 37˚C overnight, washed twice with sterile water, and allowed to dry before use. Cortices were dissected in ice-cold HBSS, and the isolated brain tissue was minced and digested with papain (Worthington Biochemical) at 20 U $mL^{-1}$ for 25 min at 37˚C. Digestion was quenched with 10% FBS in HBSS, followed by gentle trituration. Cells were pelleted at 300 × g for 7 min, resuspended in Neurobasal Plus medium (Thermo Fisher Scientific) supplemented with 1× B27 Plus (Thermo Fisher Scientific) and 1× penicillin-streptomycin[3], and seeded at $1.5 \times 10^5$ cells per culture in 150 µL medium. Half-medium changes were performed at DIV3, DIV6, and DIV9.

#### Lentivirus production and neuronal transduction

Lentivirus production and handling were performed in accordance with institutional biosafety guidelines under a protocol approved by the Institutional Biosafety Committee of the California Institute of Technology. Lentivirus encoding MagLOV was produced in HEK293T cells. Approximately $6 \times 10^6$ HEK293T cells were seeded in each 10-cm dish 24 h before transfection, with five dishes used for virus production. For each dish, 22 µg pLenti-EF1s-MagLOV-WPRE, 22 µg pKging packaging plasmid, and 4.5 µg pVSV-G, the latter two available in the laboratory, were cotransfected using linear PEI (Polysciences) at a PEI:DNA mass ratio of 2.58:1. At 12 h after transfection, the medium was replaced with complete DMEM supplemented with sodium butyrate (10 mM), followed by replacement with fresh complete DMEM 8 h later. Viral supernatant was collected after an additional 48 h, clarified by centrifugation at 500 × g for 10 min, and mixed with Lenti-X Concentrator (Takara) at 3:1 supernatant:Lenti-X volume ratio. After overnight incubation at 4˚C, virus was aliquoted and stored at -80˚C until use. For neuronal transduction, 100 µL concentrated MagLOV lentivirus was added to each neuronal culture at DIV4, and cells were imaged at DIV11.

**Magnetic field stimulation and fluorescence imaging**

Fluorescence imaging was performed at 37˚C on a custom heating stage, consisting of a circular aluminum block machined in-house with a circular well sized to hold a 35-mm dish, heated under thermocouple-based feedback control (TC-324C, Warner Instruments), on an upright microscope equipped with a 10× objective (Leica), a 470-nm LED light source (Lumencor), and an sCMOS camera (Zyla 5.5, Andor). Excitation intensity was controlled via software and calibrated using a power meter (Thorlabs). Images were acquired at 2 frames $s^{-1}$ for 60 s. Magnetic field stimulation was applied using a permanent magnet (K&J Magnetics, B666) mounted on a servo motor (Hiwonder, 20 kg cm digital servo with waterproof metal gearing; control range 180˚), which alternated the magnet between 0˚ and 90˚ positions at 10-s intervals to generate field-OFF and field-ON states, respectively, producing three magnetic field-ON periods during each acquisition. Before each experiment, the magnet was positioned such that its center was aligned with the optical axis of the excitation beam, and its height was adjusted until the magnetic field strength at the sample plane, measured using a gaussmeter (F.W. Bell), reached the target value for that experiment; the corresponding field-OFF position was confirmed by gaussmeter to read near-zero field. Sham controls underwent identical servo movement in the absence of the magnet.

**Excitation-intensity and magnetic field-strength measurements**

To characterize the dependence of MagLOV magnetofluorescence on optical excitation, cells were imaged under 60 mT magnetic field stimulation at excitation intensities of 1.65, 4.95, 9.9, 19.8, 41.2, and 82.5 mW $cm^{-2}$. To characterize the magnetic field-strength dependence, cells were imaged at a fixed excitation of 9.9 mW $cm^{-2}$ while the magnetic field strength was varied between 2, 4, 8, 15, 30, and 60 mT. All other imaging parameters were as described above.

**Fresh-medium exposure experiments**

To examine the effect of fresh-medium exposure on MagLOV magnetofluorescence, HEK293T cells were transfected as described above and imaged 48 h after transfection. Independent dishes received 3 mL fresh complete DMEM at 4, 8, 12, 16, and 20 h before a common imaging endpoint, in addition to the 500 µL culture medium already present in each dish. All conditions were imaged using an excitation intensity of 9.9 mW $cm^{-2}$ and a magnetic field strength of 4 mT.

**Fluorescence image analysis and single-cell trace extraction**

Fluorescence image stacks were analyzed on a single-cell basis using ImageJ (version 1.54p)[4]. Cell masks were generated using CellPose (version 3.1.0) with the Cyto3 model[5], applied to an average-intensity z-projection of each fluorescence image stack, with contrast adjusted prior to segmentation for the purpose of mask generation only; following manual calibration of the expected cell diameter, the resulting masks were saved as regions of interest (ROIs) and applied without further manual curation. For each identified cell, the mean fluorescence intensity within the corresponding mask was measured at every frame to generate a 120-frame fluorescence time course. The same cell mask was applied throughout each acquisition. Single-cell traces were subsequently processed in Python (version 3.11.5) for fluorescence correction and quantitative analysis as described below.

**Fluorescence drift correction**

Single-cell fluorescence traces were corrected for slow illumination-associated drift and an initial fluorescence transient using a log-space regression model,

$$\log F(t) = a + b_1 t + b_2 t^2 + b_3 t^3 + Ae^{-t/\tau} + \beta S(t) \ ,$$

where $S(t)$ denotes the magnetic field state, taking a value of 0 during field-OFF intervals and 1 during field-ON intervals. Cells with any non-positive or missing fluorescence values were excluded prior to fitting. Fits were performed using frames 1–18, 29–38, 49–58, 69–78, 89–98, and 109–118; the first interval spans the full initial OFF period, including the startup transient to be modeled, whereas subsequent intervals use only the final 10 frames of each 20-frame OFF or ON block to avoid field-switching kinetics. The cubic baseline and exponential startup components were treated as nuisance terms and removed, whereas the field-state term was retained. For each biological replicate, the shared startup time constant $\tau$ was estimated by a grid search over 191 candidate values spanning 0.5–10 s (spacing ≈ 0.05 s); for each candidate value, the remaining coefficients were solved by ordinary least squares (NumPy 1.24.3, linalg.lstsq) using the population-average log-fluorescence trace across all cells in that replicate, and the value minimizing the residual sum of squares was selected. The fitted $\tau$ did not approach either search boundary in any condition. This shared $\tau$ was then used to fit the remaining coefficients independently for each cell within the replicate by ordinary least squares. Corrected log-fluorescence values were exponentiated and expressed as relative fluorescence, with $\Delta F/F_0 = (F_{\mathrm{rel}} - 1) \times 100\%$.

**MFE amplitude and response-kinetics analysis**

Plateau MFE amplitudes were calculated from drift-corrected single-cell fluorescence traces as

$$\mathrm{MFE} = \frac{F_{\mathrm{OFF}} - F_{\mathrm{ON}}}{F_{\mathrm{OFF}}} \times 100\% \ ,$$

where $F_{\mathrm{OFF}}$ was the mean fluorescence during the final 2 s preceding magnetic field onset and $F_{\mathrm{ON}}$ was the mean fluorescence during the final 2 s of the corresponding 10-s field-ON interval; these 2-s windows were nested within the larger windows used for drift-model fitting (see above), capturing the portion of each interval closest to steady state. MFE amplitudes from the three stimulation cycles were first averaged within each cell and then across cells within each biological replicate. Cells with any non-finite amplitude value in any cycle were excluded prior to averaging. Response kinetics were quantified from drift-corrected biological-replicate mean traces. For each OFF-to-ON transition, the response was normalized as

$$R(t) = \frac{F_{\mathrm{baseline}} - F(t)}{F_{\mathrm{baseline}} - F_{\mathrm{plateau}}} \ ,$$

where $F_{\mathrm{baseline}}$ and $F_{\mathrm{plateau}}$ were the mean fluorescence during the pre-field and final field-ON windows defined above, such that $R = 0$ at baseline and $R = 1$ at plateau. Cycles in which the field-associated fluorescence change was not positive ( $F_{\mathrm{baseline}} - F_{\mathrm{plateau}} \leq 0$ ) were excluded from kinetic analysis. The response half-time ($t_{1/2}$) was defined as the first time point, searched from the onset of the field-ON interval through the end of the plateau window, at which $R(t)$ reached 0.5, determined by linear interpolation between adjacent frames; if $R(t)$ at the first field-ON frame already exceeded 0.5, $t_{1/2}$ was set to 0. Cycle-specific $t_{1/2}$ values were averaged within each biological replicate, using only cycles that met the inclusion criterion above.

**Correction-independent robustness analyses**

To verify that the measured magnetofluorescence was not dependent on the drift-correction procedure, additional analyses were performed directly on raw fluorescence traces. For each magnetic field transition, the response was calculated as $(F_{\mathrm{OFF}} - F_{\mathrm{ON}})/F_{\mathrm{OFF}} \times 100\%$, where

$F_{\mathrm{OFF}}$ and $F_{\mathrm{ON}}$ denote the mean fluorescence during OFF- and ON- state windows, respectively, regardless of their temporal order. $F_{\mathrm{OFF}}$ corresponds to the pre-switch window for OFF-to-ON transitions and to the post-switch window for ON-to-OFF transitions. For both transition directions, the pre-switch window comprised the 2 s (4 frames) immediately preceding the switch, and the post-switch window comprised a 2-s window beginning 2 s after the switch, omitting the intervening 2-s period to avoid switching kinetics. This yielded three OFF-to-ON and two ON-to-OFF transitions per acquisition, reflecting the number of complete transitions of each type captured within the 60-s recording. Transition-specific values of the same direction were first averaged within each cell and then across cells within each biological replicate. A direction-balanced estimate was calculated for each biological replicate as the unweighted mean of the OFF-to-ON and ON-to-OFF estimates. Robustness to the choice of post-switch analysis window was assessed by repeating this analysis with the post-switch window beginning 0, 1, 2, or 3 s after each transition, with the pre-switch window held fixed. No drift, startup-transient, or phase-dependent correction was applied in these analyses.

**Basal fluorescence and correlation analyses**

Basal MagLOV fluorescence was quantified from uncorrected fluorescence traces as the mean intensity over frames 1–18 for each cell and subsequently averaged across cells within each biological replicate. For relative comparisons across fresh-medium exposure times (Extended Data Fig. 3c), replicate-level basal fluorescence and MFE amplitude were independently normalized to the corresponding matched 4-h value. Independently of this normalization, the relationship between raw (non-normalized) basal fluorescence and MFE amplitude was assessed across all biological replicates using two-tailed Pearson correlation and ordinary least-squares linear regression with a free intercept and 95% confidence bands (Extended Data Fig. 3d). To assess this relationship independently of exposure-time-dependent group shifts, raw basal fluorescence and MFE amplitude were centered within each fresh-medium exposure group by subtracting the corresponding group mean, and the centered values were pooled for Pearson correlation analysis (Extended Data Fig. 3e).

**Biological-replicate and cycle-level robustness analyses**

To assess the reproducibility of MagLOV responses across independent biological replicates, raw single-cell fluorescence traces were normalized to the mean fluorescence over frames 1–18 for each cell and then averaged across cells within each biological replicate; cells with a non-finite or non-positive baseline value were excluded. No drift, startup-transient, or phase-dependent correction was applied to these traces. To assess response stability across repeated magnetic field stimulation, plateau MFE amplitudes were calculated separately for each of the three field-ON cycles from drift-corrected single-cell traces using the definition described above; cells with a non-finite amplitude value in any cycle were excluded. Cycle-specific amplitudes were averaged across cells within each biological replicate, and the same biological replicates were compared across successive stimulation cycles (see Statistical analysis).

**Statistical analysis**

Statistical analyses were performed using GraphPad Prism (version 11.0.1). Independent biological replicates, rather than individual cells, were treated as the statistical unit. Unless otherwise indicated, summary data are presented as mean ± s.d., whereas fluorescence time courses are shown as mean ± s.e.m. Comparisons among multiple independent conditions were performed using one-way ANOVA followed by Dunnett's multiple-comparisons test using the indicated reference condition. Cycle-resolved measurements from the same biological replicates were analyzed by repeated-measures one-way ANOVA with Geisser-Greenhouse correction followed by Dunnett's multiple-comparisons test against Cycle 1. Associations between continuous variables were assessed using two-tailed Pearson correlation and ordinary least-squares linear regression with a free intercept. Statistical significance was defined as $P < 0.05$. Exact sample sizes and statistical tests are specified in the corresponding figure legends.

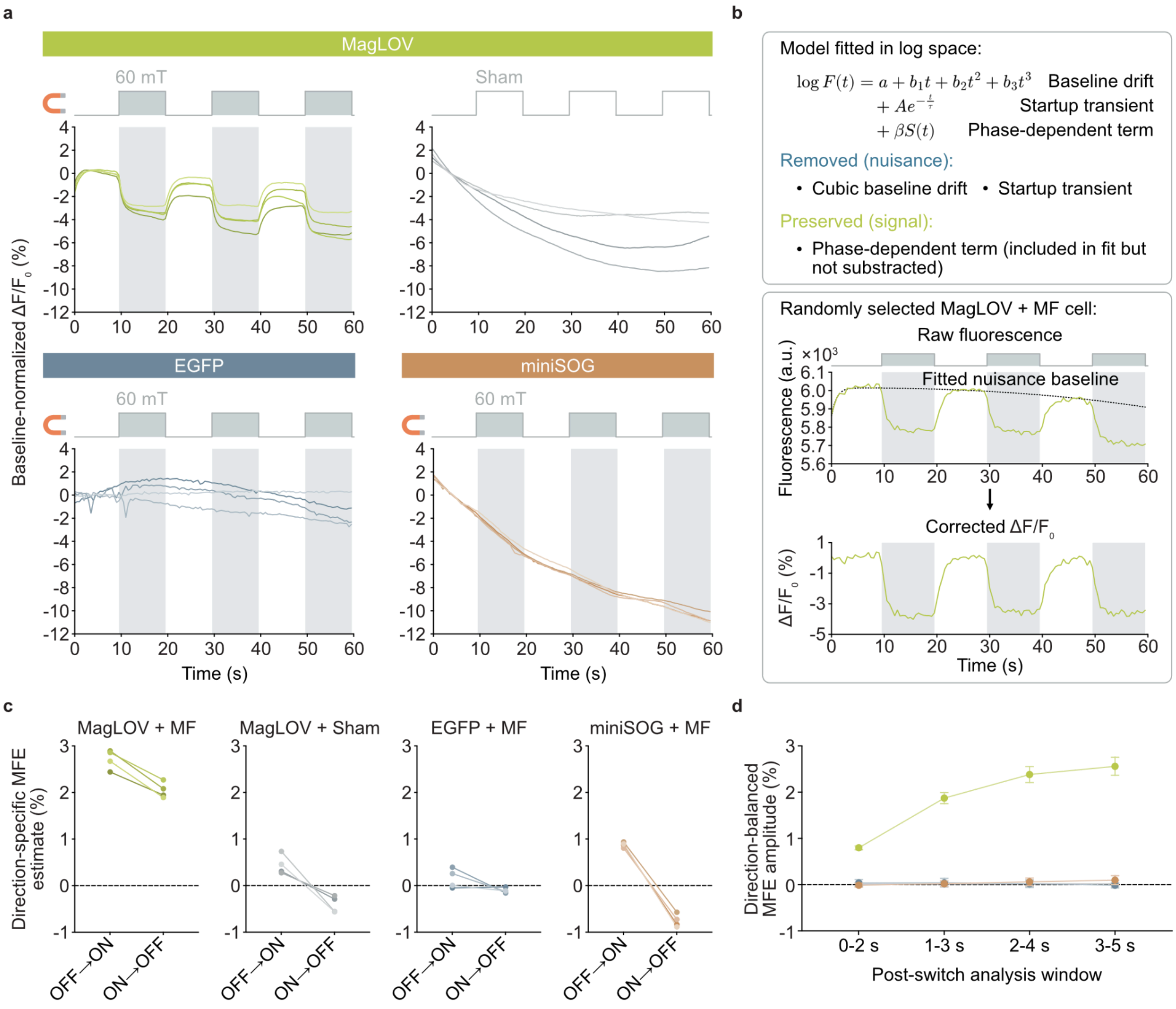


**Extended Data Fig. 1 | Validation of fluorescence correction and robustness of MagLOV MFE detection**

**a,** Baseline-normalized fluorescence traces from individual biological replicates before drift correction for HEK293T cells expressing MagLOV during 60 mT magnetic field stimulation or sham servo movement, and cells expressing EGFP or miniSOG during 60 mT stimulation. For each cell, fluorescence was normalized to the mean fluorescence before the first field switch (see Methods). Each line represents one independent biological replicate (n = 4 per condition). Shaded regions indicate magnetic field-on intervals; outlined intervals indicate the corresponding servo phases for the sham condition.

**b,** Fluorescence-correction procedure (see Methods). Top, decomposition of a fluorescence trace into nuisance (baseline drift, startup transient) and signal (phase-dependent) components. Bottom, representative raw fluorescence trace from a randomly selected MagLOV + MF cell, the fitted nuisance baseline, and the resulting corrected $\Delta F/F_0$ trace.

**c,** Direction-specific MFE estimates calculated directly from raw fluorescence for OFF-to-ON and ON-to-OFF magnetic field transitions (see Methods). Estimates were oriented such that a magnetic field-associated fluorescence decrease corresponds to a positive MFE in both switching directions. Connected points represent the same independent biological replicate (n = 4 per condition). The dashed line indicates zero MFE.

**d,** Robustness of the correction-independent, direction-balanced MFE estimate to the post-switch analysis window (see Methods). This analysis was performed directly on raw fluorescence traces and is independent of the plateau-amplitude quantification used in Fig. 1e. Points and error bars indicate the mean ± s.d. across four independent biological replicates. The dashed horizontal line indicates zero MFE.

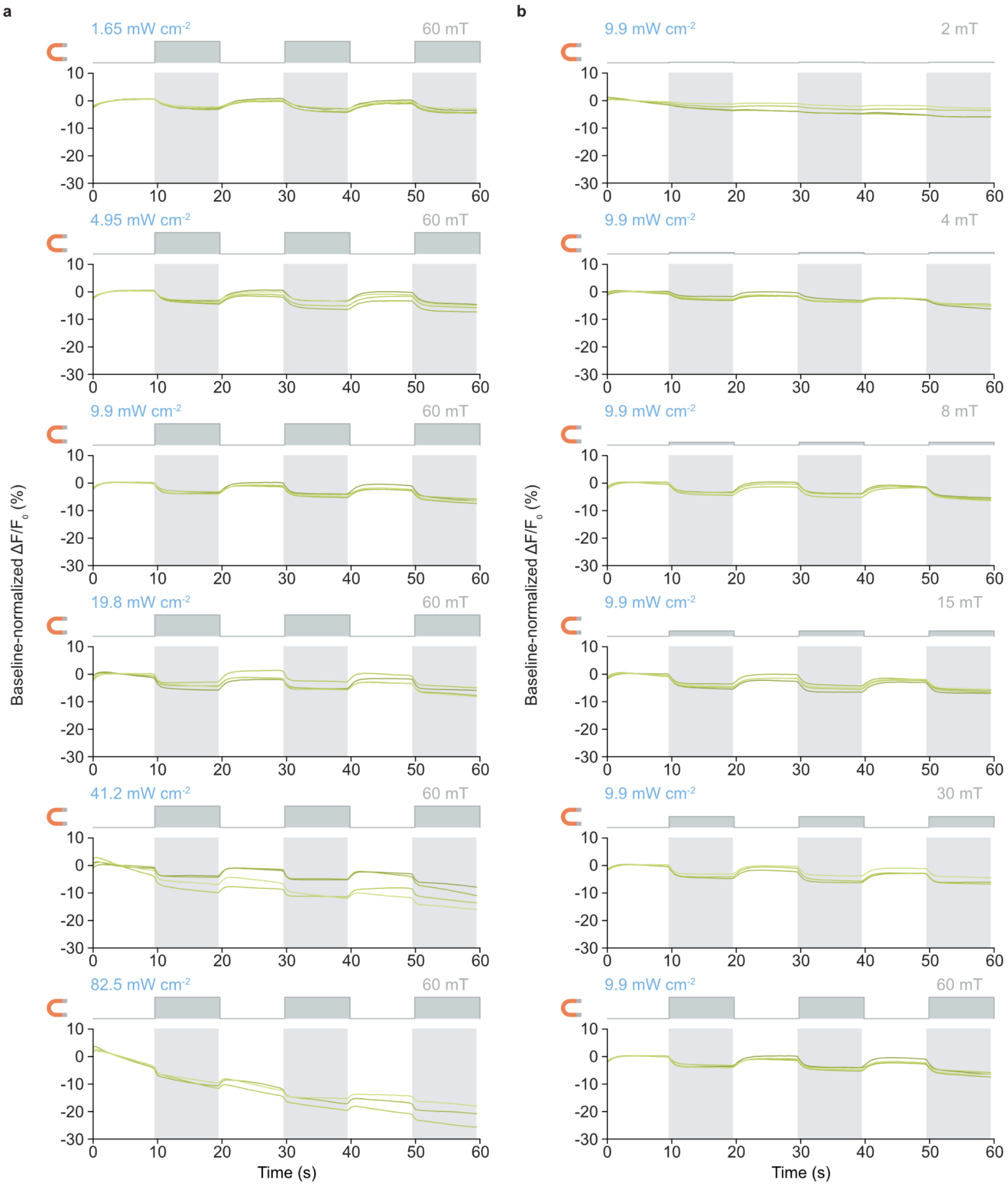


**Extended Data Fig. 2 | Uncorrected biological-replicate responses across excitation intensities and magnetic field strengths**

**a,** Baseline-normalized fluorescence traces from individual biological replicates of MagLOV-expressing HEK293T cells during 60 mT magnetic field stimulation at the indicated excitation intensities (see Methods). Each line represents one independent biological replicate.

**b,** Baseline-normalized fluorescence traces from individual biological replicates of MagLOV-expressing HEK293T cells during stimulation with the indicated magnetic field strengths at a fixed excitation intensity of 9.9 mW cm$^{-2}$. Each line represents one independent biological replicate. **a,b,** Shaded regions indicate magnetic field-on intervals. No drift, startup-transient or phase-dependent correction was applied. n = 3–4 independent biological replicates per condition.

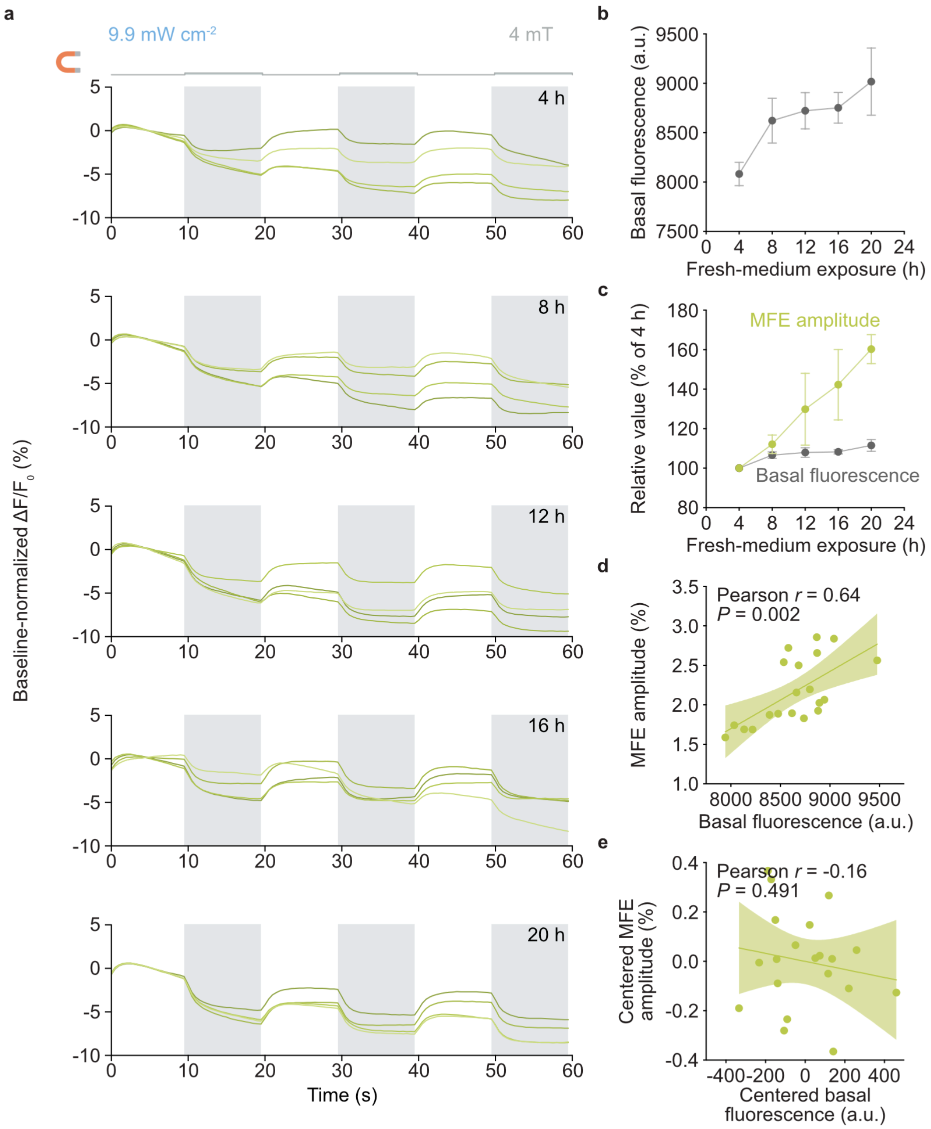


**Extended Data Fig. 3 | Uncorrected biological-replicate responses and basal fluorescence correlates of the fresh-medium exposure effect**

**a,** Baseline-normalized fluorescence traces from individual biological replicates of MagLOV-expressing HEK293T cells during 4-mT magnetic field stimulation at an excitation intensity of 9.9 mW $cm^{-2}$, following the indicated durations of fresh-medium exposure. Each line represents one independent biological replicate (n = 4 per condition). Shaded regions indicate magnetic field-on intervals. No drift, startup-transient or phase-dependent correction was applied.

**b,** Basal MagLOV fluorescence as a function of fresh-medium exposure duration, quantified from uncorrected fluorescence traces (see Methods). Points and error bars indicate the mean ± s.d. across four independent biological replicates (n = 4).

**c,** Basal fluorescence and MFE amplitude from **b** and Fig. 3c, respectively, each normalized to their matched 4-h value (see Methods). Points and error bars indicate the mean ± s.d. across four independent biological replicates (n = 4).

**d,** Relationship between basal fluorescence and MFE amplitude across all biological replicates and fresh-medium exposure times. Line and shaded band indicate the ordinary least-squares linear fit and 95% confidence interval. Pearson correlation coefficient and P value are indicated.

**e,** As in **d**, after centering basal fluorescence and MFE amplitude within each fresh-medium exposure group to remove exposure-time-dependent group shifts (see Methods). Pearson correlation coefficient and P value are indicated.

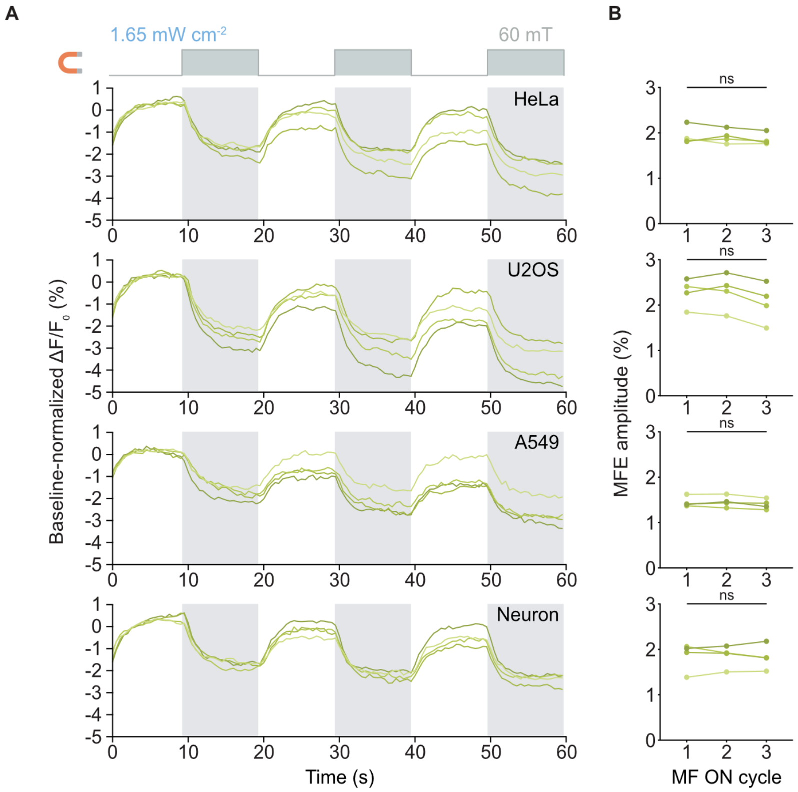


**Extended Data Fig. 4 | Replicate- and cycle-level robustness of MagLOV magnetofluorescence across diverse mammalian cellular backgrounds**

**a,** Baseline-normalized fluorescence traces from individual biological replicates of MagLOV-expressing HeLa, U2OS, and A549 cells and primary mouse cortical neurons during repeated 60 mT magnetic field stimulation at an excitation intensity of 1.65 mW $cm^{-2}$ (see Methods). Each line represents one independent biological replicate (n = 4 per cellular background). Shaded regions indicate magnetic field-on intervals. No drift, startup-transient or phase-dependent correction was applied.

**b,** Cycle-resolved plateau MFE amplitudes for HeLa, U2OS, and A549 cells and primary mouse cortical neurons, calculated using the same definition as in Fig. 4c (see Methods). Connected points represent the same independent biological replicate across successive stimulation cycles. Mean ± s.d. is shown for each cycle. Statistical significance was assessed separately within each cellular background using repeated-measures one-way ANOVA with Geisser-Greenhouse correction followed by Dunnett's multiple-comparisons test against Cycle 1; ns, not significant.
